# Translating deep learning to neuroprosthetic control

**DOI:** 10.1101/2023.04.21.537581

**Authors:** Darrel R. Deo, Francis R. Willett, Donald T. Avansino, Leigh R. Hochberg, Jaimie M. Henderson, Krishna V. Shenoy

## Abstract

Advances in deep learning have given rise to neural network models of the relationship between movement and brain activity that appear to far outperform prior approaches. Brain-computer interfaces (BCIs) that enable people with paralysis to control external devices, such as robotic arms or computer cursors, might stand to benefit greatly from these advances. We tested recurrent neural networks (RNNs) on a challenging nonlinear BCI problem: decoding continuous bimanual movement of two computer cursors. Surprisingly, we found that although RNNs appeared to perform well in offline settings, they did so by overfitting to the temporal structure of the training data and failed to generalize to real-time neuroprosthetic control. In response, we developed a method that alters the temporal structure of the training data by dilating/compressing it in time and re-ordering it, which we show helps RNNs successfully generalize to the online setting. With this method, we demonstrate that a person with paralysis can control two computer cursors simultaneously, far outperforming standard linear methods. Our results provide evidence that preventing models from overfitting to temporal structure in training data may, in principle, aid in translating deep learning advances to the BCI setting, unlocking improved performance for challenging applications.

## Introduction

Rapid progress in machine learning and artificial intelligence has led to an impressive collection of neural network models capable of learning complex nonlinear relationships between large amounts of data (these approaches have been referred to as “deep learning”). Deep learning algorithms have produced significant success in a wide variety of applications^1^ including, computer vision^2–4^, natural language processing^5–7^, and robotics^8–10^. More recently, a promising application of neural networks has been towards modeling and decoding the brain activity associated with movement via brain-computer interfaces (BCIs), which holds great potential for improving performance of BCI systems. However, this intersection of deep learning and BCIs presents some unique challenges, including the often-limited quantity of data and changes in the distribution of data from the offline (open-loop) to online (real-time closed-loop control) settings.

Intracortical BCIs are systems that aim to restore movement and communication to people with paralysis by decoding movement signals from the brain via microelectrodes placed in the cortex. Advancements in clinical research BCIs have enabled functional restoration of movement and communication, including robotic arm control^11–14^, reanimation of paralyzed limbs through electrical stimulation^15–19^, cursor control^20–22^, decoding speech^23–27^, and most recently, decoding handwriting^28^. An abundance of prior work suggests that BCI decoding may be improved through neural networks, as demonstrated in various offline settings^29–35^. To date, however, there are only a few demonstrations of continuous online BCI control using neural networks, most of which are restricted to nonhuman primate (NHP) studies^36, 37^ given the rarity of real-time human BCI data. Many prior motor decoding algorithms for real-time neuroprosthetic control – which convert movement-related brain activity into continuous control signals – have been based on linear methods^13, 38–43^. Here, we apply a neural network for real-time neuroprosthetic control to assess whether it can generate advances in performance as suggested by prior work.

Of the many network architectures, recurrent neural networks (RNNs) have been a popular decoding approach for BCIs^29, 33, 36^ since they can learn temporal dependence within data, aligned with the dynamical systems view that neural activity in the motor cortex evolves over time^29, 44, 45^. However, RNNs often require large amounts of training data and can overfit to the temporal structure within offline data which may not be present in online data, potentially reducing their utility as decoders for BCI applications. In this study, we investigate the usage and application of RNNs on a challenging nonlinear BCI problem: controlling two cursors simultaneously via decoded bimanual movement.

Prior studies have shown that motor cortex contributes to both contralateral and ipsilateral movements and that neural tuning changes nonlinearly between single and dual-limb movements^46–51^. More specifically, during dual movement we found that the neural representation for one effector (‘primary’) stays relatively constant, whereas the other effector’s (‘secondary’) representation gets suppressed while its directional tuning changes. Additionally, there is significant correlation in how movement direction is represented for contralateral and ipsilateral movements. To date, studies that have investigated bimanual BCI control^41, 46, 52, 53^ have mainly used linear decoding algorithms (e.g., Kalman filters and ridge regression) despite the seemingly nonlinear relationship between neural activity and bimanual movement. The need for exploration of nonlinear decoding methods for bimanual movement makes this problem an apt application and testbed for RNNs.

Here, we demonstrate a surprising finding: that RNN decoders calibrated on stereotyped training data achieve high offline performance (consistent with prior work^33, 54–56)^, but do so in part by overfitting to the temporal structure of the task, resulting in poor performance when used for online, real-time control of a BCI. To solve this problem, we altered the stereotyped structure in training data to introduce temporal and behavioral variability which helps RNNs generalize to the online setting. In addition, we show that RNNs can leverage nonlinearities within the neural code governing complex bimanual movements to accomplish simultaneous two-cursor control, outperforming linear methods. Overall, our findings suggest that preventing overfitting to temporal structure within training data can help translate advances in deep learning to improve BCI performance on challenging nonlinear problems.

## Results

### Nonlinear neural coding of directional unimanual and bimanual hand movement

We first sought to understand how bimanual hand movements are represented in motor cortex, including sources of nonlinearity that would motivate the use of RNNs. We used microelectrode recordings from the hand knob area of the left (dominant) precentral gyrus in a clinical trial participant (referred to as T5) to characterize how neural tuning changes between bimanual hand movement (both hands attempting to move simultaneously) and unimanual hand movement (one hand moving individually). T5 has a C4 spinal cord injury and is paralyzed from the neck down; attempted movement resulted in little to no motion of the arms and legs (see Willett*, Deo*, et al. 2020 for more details^50^). T5 was instructed to attempt hand movements.

Using a delayed movement task (Fig. 1a), we measured T5’s neural modulation to attempted unimanual and bimanual hand movements. We observed changes in neural spiking activity across many individual electrodes as a function of movement direction during bimanual movements (Fig. 1b presents an example electrode’s responses; see Supplementary Fig. 1 for a count of tuned electrodes). We also observed nonlinear changes in tuning from the unimanual to bimanual context, including tuning suppression and direction changes (Fig. 1c). Here, ‘nonlinear’ is considered any departure from linear tuning to the variables we intend to decode: the x- and y-components of movement direction^41^, as described in the encoding model (equation 1) below:

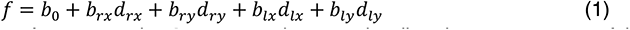

**Fig. 1.**
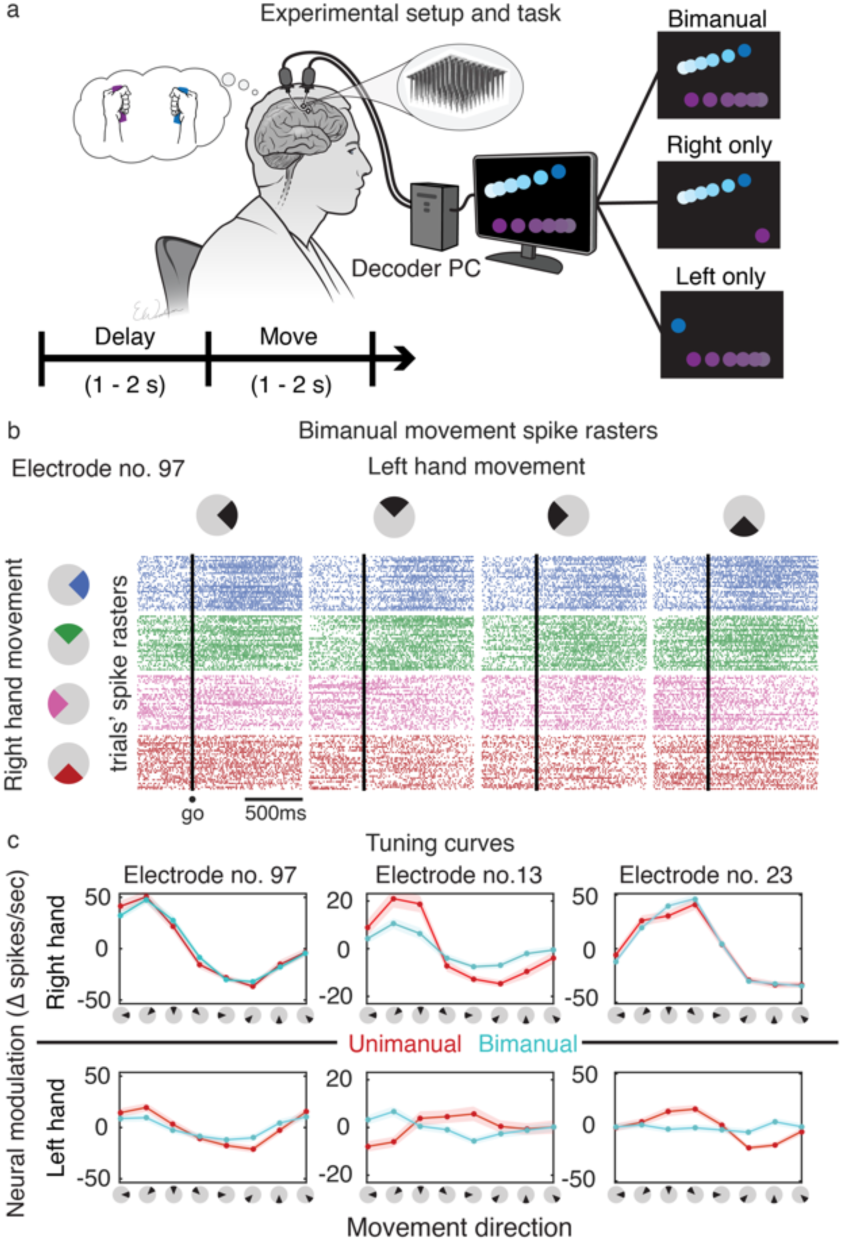
Neural tuning to unimanual and bimanual hand movement. **a** Participant T5 performed a delayed-movement task. Cursors on a screen prompted T5 to attempt to make concomitant joystick movements. One of three types of movements were cued on each trial: (1) *bimanual*: both hands, (2) *unimanual left*: only left (ipsilateral) hand, or (3) *unimanual right*: only right (contralateral) hand. **b** Matrix of spike rasters of example electrode no. 97 during bimanual movements. Raster plot (i,j) of the matrix corresponds to electrode 97’s response to right hand movement in direction *i* while the left hand moved in direction *j* (colored by right hand direction). Each row of a raster plot represents a trial and each column is a millisecond time-bin. A dot indicates a threshold crossing spike at the corresponding trial’s time-bin. Different spiking activity can be seen for different bimanual movements, indicating tuning to bimanual movement direction. **c** Tuning curves of example electrodes show a range of tuning changes to each hand (rows) across movement contexts (red/blue). Solid dots indicate the mean firing rates (zero-centered) for movements in the directions indicated on the x-axes. Spikes were binned (20-ms bins) and averaged within a 300-700 ms window after the ‘go’ cue. Shaded areas are 95% CIs (computed via bootstrap resampling). Electrode no. 97 retained tuning for both hands between contexts, electrode no. 13 had suppressed tuning for both hands during bimanual movement, and electrode no. 23 had suppression in left hand tuning during bimanual movement.

Here, *f* is the average firing rate of a neuron, the d terms are the *x*- and *y*-direction components of the right (*d_rx_*, *d_ry_*) and left (*d_lx_*, *d_ly_*) hand velocities, and the *b* terms are the corresponding coefficients of the velocity components (and *b_0_* is the baseline firing rate). Tuning angle changes (“decorrelation”) and a suppressed tuning magnitude from unimanual to bimanual movement breaks linearity, since the tuning coefficients change based on movement context. In addition, direction-independent laterality tuning (i.e., coding for the side of the body irrespective of movement direction) is another potential key source of nonlinearity. For clarity, Figure 2a illustrates these three nonlinear phenomena (decorrelation, suppression, and laterality tuning) with a schematic.

**Fig. 2.**
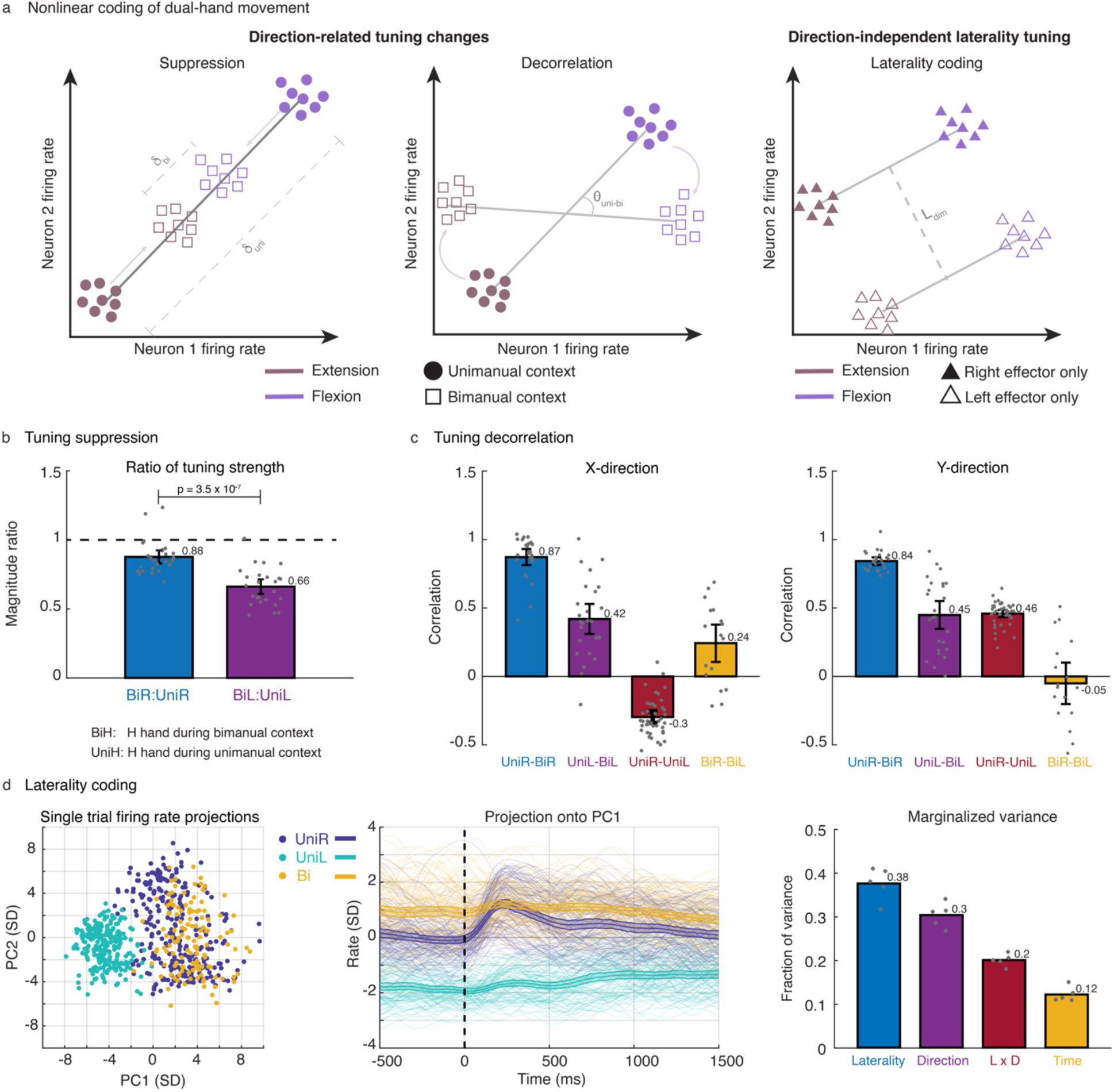
Nonlinear neural code underlying bimanual hand movement. **a** Cartoon examples of three key sources of nonlinearity in the neural coding of directional bimanual and unimanual movement. Firing rates for two exemplary neurons are plotted for flexion (purple) and extension (brown) of an effector (left and middle panels) during unimanual and bimanual contexts, or for two effectors (right panel) during unimanual movement. Direction-related tuning changes consist of suppression (reduction in neural distance between movement representations), and decorrelation (change in tuning axis) between unimanual and bimanual contexts. Direction-independent laterality tuning can be viewed as a large dimension separating movements between effectors on opposite sides of the body. **b** Amount of tuning suppression in offline data. Each bar indicates the mean ratio of tuning strength between bimanual and unimanual contexts for right (blue) and left (purple) hand movement. Significance was determined by a 2-sample t-test. All black intervals on bar plots indicate 95% CIs. Left hand tuning was suppressed more during the bimanual context than right hand tuning. **c** Degree of tuning decorrelation in offline data. Each bar indicates the correlation between the neural population’s x- or y-direction coefficient vectors for pairs of movement types. See Supplementary Table 1 for p values. Right hand directional tuning remained largely unchanged while left hand directional tuning changed more substantially during the bimanual context. **d** Laterality information in offline data. Principal component analysis (PCA) on single trial Z-scored firing rates (SD denotes standard deviation) is used to visualize how movement types cluster (left panels; each dot and line is a single trial). Demixed PCA was used to compute the marginalized variance of different movement factors (right panel). Tuning to laterality was stronger than tuning to movement direction.

Tuning decorrelation and a suppression of ipsilateral related neural activity have been seen previously during bimanual movement^47, 50^. These phenomena can be reproduced even with a richer set of continuous directional movements (Fig. 2b). The tuning strength of right hand (primary effector) directional movements remained relatively unchanged from unimanual to bimanual contexts (12% suppression during bimanual), whereas tuning strength of the left hand (secondary effector) was suppressed by 34% during bimanual movement. Similarly, directional tuning (Fig. 2c) of the right hand remained relatively unchanged (0.87 and 0.84 correlations for x- and y-directions, respectively) while left hand directional tuning changed more substantially (0.42 and 0.45 correlations for x- and y-directions, respectively) from the unimanual to bimanual context. These results indicate that neural tuning to left hand movements exhibited suppression and decorrelation when moved simultaneously with the right hand, whereas tuning to right hand movements remained mostly unchanged.

### A large neural dimension codes for laterality of the hand

Also consistent with our prior work, we found a salient laterality-related neural dimension (Fig. 2d) coding for the side of the body that the hand resides independent of the movement direction. We used principal component analysis (PCA) on both unimanual and bimanual neural data to visualize neural activity in the top principal components (PCs). A dimension emerged within the top two PCs clearly separating right from left hand unimanual movements. Interestingly, bimanual neural activity most closely resembled that of unimanual right hand activity in the top PCs, further indicating that the right hand is more strongly represented than the left hand during bimanual movement in the contralateral precentral gyrus. Next, we used demixed PCA^57^ (dPCA), which decomposes neural data into a set of dimensions that each explain variance related to one marginalization of the data, to quantify the size of the laterality factor in unimanual movement data only. We marginalized the data according to the following factors: time, laterality, movement direction, and the laterality-direction interaction. The laterality marginalization contained the highest fraction of variance (39% marginalized variance) indicating that tuning to laterality was stronger than tuning to direction (30% marginalized variance). From a decoding perspective, laterality dimensions can be useful in distinguishing right hand movements from left hand movements in a unimanual context.

Overall, we found a strong presence of nonlinearities within the neural code governing bimanual hand movement, making this a well suited application for RNN decoding.

### RNNs overfit to the temporal structure of offline data and generate overly stereotyped online behavior

Next, we used a simple RNN architecture (Fig. 3a and Supplementary Fig. 2) – similar to the neural network model used in our recent report on decoding attempted handwriting^28^ – to decode bimanual movement from neural activity. During RNN calibration, neural activity was recorded while T5 attempted movements in concert with one or both cursors moving on a screen. The structure of this task followed a delayed movement paradigm where T5 *prepared* to move during a delay period, executed movement during a *move* period, and then rested at an *idle* state. This highly stereotyped temporal structure (*prepare-move-idle*) is typical of BCI calibration tasks in which neural activity can be regressed against the inferred behavior. The RNN was trained to convert neural activity into (1) left and right cursor velocities and (2) discrete movement- context signals that denoted the category of movement being made at each moment in time (unimanual left, unimanual right, bimanual, or no movement). During closed-loop cursor control, the discrete context signals were used to gate the output cursor velocities. Velocity targets for RNN training were modified by introducing a reaction time and saturating the velocity curve (Fig. 3b) to better approximate the participant’s intention to move maximally when far from the target^58^.

**Fig. 3.**
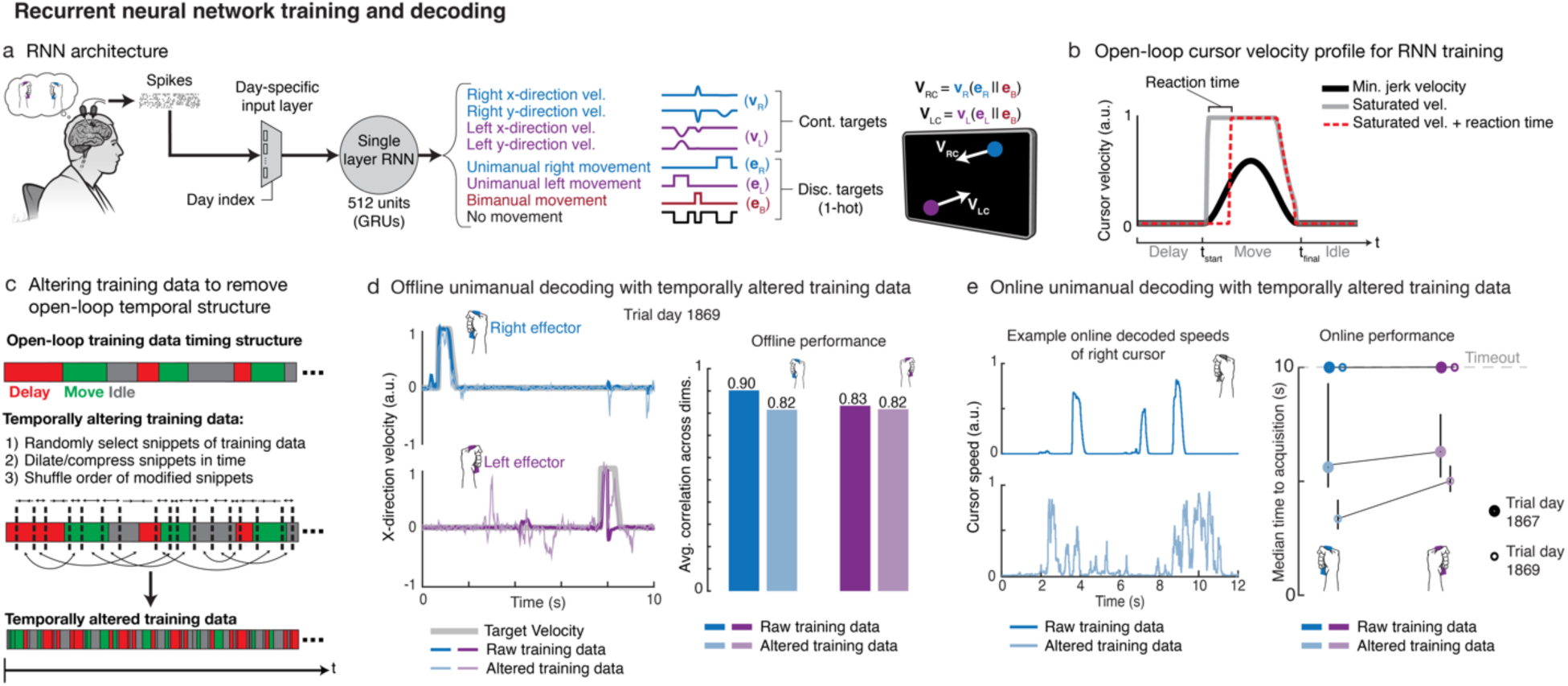
Fracturing temporal structure in offline training data helps RNN decoders generalize to the online setting. **a** Diagram of the decoding pipeline. First, neural activity (multiunit threshold crossings) was binned on each electrode (20-ms bins). Then, a trainable day-specific linear input layer transformed the binned activity from a specific day into a common space to account for day-to-day variabilities in signal recordings. Next, an RNN converted the day- transformed time series activity into continuous left and right cursor velocities (*ν_R_*, *ν_L_*), and discrete movement context signals (*e_R_*, *e_L_*, *e_B_*). The movement context signals were then used to gate the appropriate cursor velocity outputs. **b** Example open-loop minimum-jerk cursor velocity (black) and modified saturated velocities (gray/red). Saturated velocity with a prescribed reaction time of 200 ms (red) is used for RNN training since it better approximates behavior. **c** Data alteration technique that introduces variability in the temporal and behavioral structure of training data. Data is subdivided into small snippets of variable length, each snippet is then dilated or compressed in time, and the order of the modified snippets are shuffled. This allows for synthetic data generation as well. **d** Offline decoding performance of RNNs trained with and without data alteration. Sample snippets of x-direction decoded velocities are shown for both cursors during unimanual movement with RNNs trained with and without alteration. Corresponding decoding performance (Pearson correlation coefficient) is summarized via bar plots. Offline performance is better without data alteration, mainly due to overfitting. **e** Decoders trained with unaltered data generated pulse-like movements online, as shown in the sample decoded cursor speeds for the right hand (top panel), whereas the RNN trained with altered data (bottom panel) allowed for quicker online corrections. Decoders trained with altered data acquired targets more quickly online.

To investigate the RNN’s decoding efficacy, we first focused on the unimanual movement case, which mitigates decoding challenges due to suppressed left-hand representation during bimanual movement. RNNs trained on open-loop unimanual movements achieved high offline decoding performance for both hands (Fig. 3d; average correlation of 0.9 and 0.83 for the right and left hand, respectively). Surprisingly, however, these RNNs generated pulse-like movements reflecting the velocity profiles used for offline training, making subsequent closed-loop online control difficult (Fig. 3e). Instead of being able to smoothly correct for inevitable errors that occur during online control, T5 had to make repeated attempted movements – mimicking the *prepare-move-idle* offline behavior – in succession to successfully acquire targets. In this scenario, offline RNN decoding on held-out test data yielded deceptively high performance which did not translate to high online performance.

### Fracturing the stereotyped temporal structure of open-loop training data helps RNNs transfer to online control

Since the RNN decoders overfit to the stereotyped *prepare-move-idle* open-loop behavior, we hypothesized that introducing variability in the temporal and behavioral structure of the training data would help generalize to the closed-loop context. To accomplish this, we developed a simple method whereby we alter the training data by randomly selecting snippets of data (ranging between 200-800ms in duration), stretching or compressing the snippets in time using linear interpolation, and then shuffling the order of the modified snippets (Fig. 3c; see Methods). This approach aims to intermix variable size windows of neural activity across the various stages of behavior (*prepare*, *move*, and *idle*) to make the RNN decoder more robust to the rapid changes in movement direction that occur during closed-loop control. Comparing the RNN trained with temporally altered data (*altRNN)* to that trained with raw data (*rawRNN)* as described in the previous section, the *altRNN* did not overfit to the open-loop task structure, which resulted in slightly poorer decoding performance on offline held-out test data and the decoded output velocities appeared generally noisier (Fig. 3d). However, the *altRNN* led to improved closed-loop control (see Supplementary Movie 4). The decoded cursor speeds were more continuous in nature and did not overfit to the pulse-like velocity profiles prescribed to the cursors during the open-loop task (Fig. 3e).

In addition to enforcing that the RNN generalizes to data with less stereotyped structure, this data alteration technique allows for synthetic data generation which also helps to prevent overfitting to the limited amount of data that can be collected in human BCI research. Overall, we found that fracturing temporal and behavioral structure in the training data resulted in more continuous output velocities which translated to better closed-loop cursor control performance.

### RNN decoders enable online simultaneous control of two cursors

Next, we tested whether an RNN decoder trained with temporally altered data could facilitate real-time neural control of two cursors at the same time – a challenging nonlinear decoding problem. To do so, we trained an RNN on offline and online unimanual and bimanual hand movements collected over multiple sessions (see Methods). T5 attempted a series of unimanual or bimanual hand movements to drive two cursors to their intended targets. To acquire targets, the cursors had to dwell within their corresponding target for 500 ms, simultaneously. T5 was asked to attempt all bimanual trials with simultaneous hand movements (as opposed to sequential unimanual movement of one cursor at a time). T5 successfully achieved bimanual control across many sessions (see Supplementary Movie 1), where time-to-acquisition (TTA) for bimanual trials was only slightly longer than the TTA for unimanual trials on average (Fig. 4a). Amongst unimanual trials, the average TTA for right- and left-hand trials was similar. The average angular errors for both hands were generally higher during bimanual movement than during unimanual movement.

**Fig. 4.**
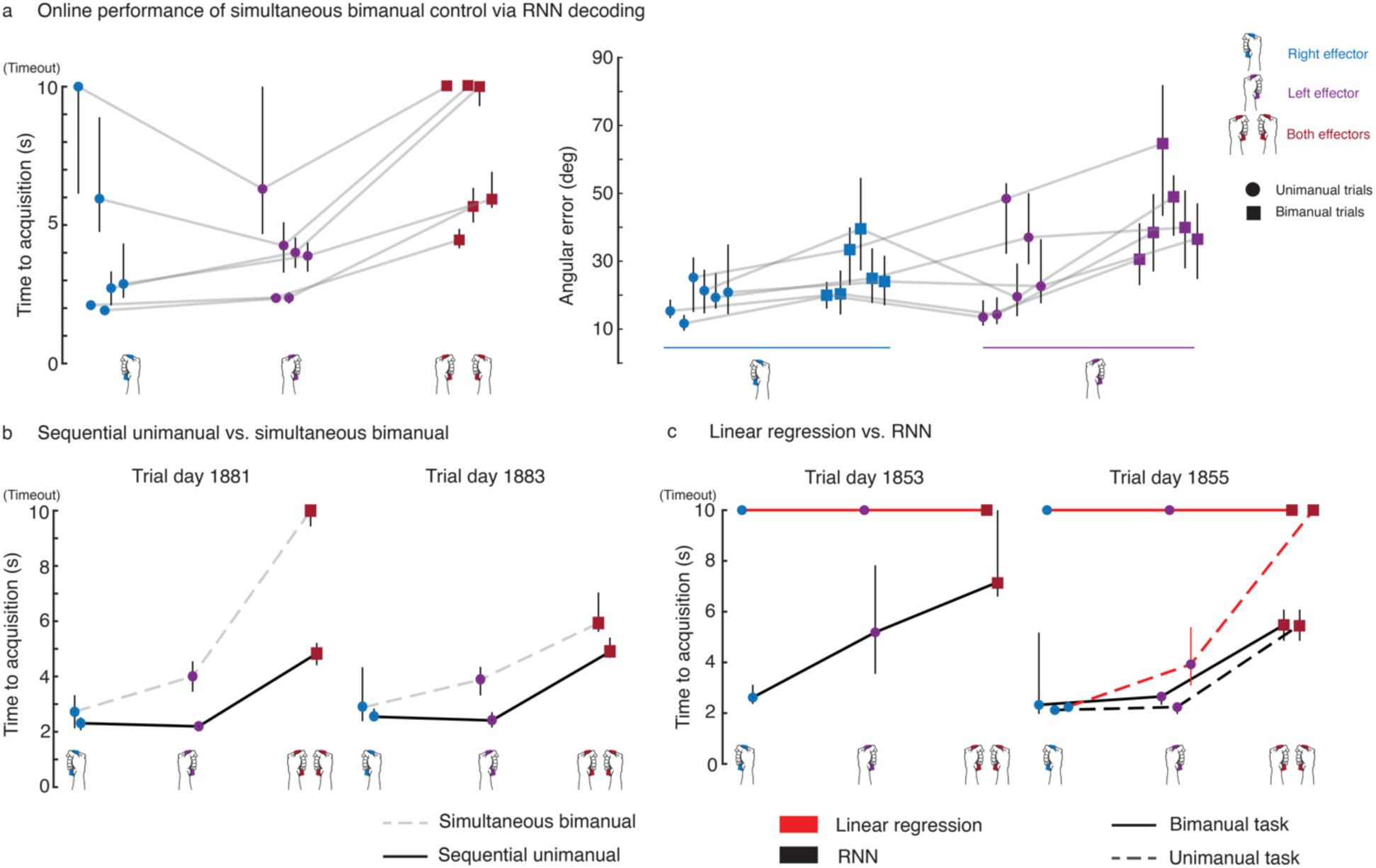
RNN decoders enable two-cursor control and outperform linear decoders. **a** Median target acquisition time and angular errors are shown for 6 days of simultaneous bimanual two-cursor control as enabled by RNN decoders. Light gray lines connect data points corresponding to the same session day. Each trial had a 10 s timeout after which the trial was considered failed. Angular error was calculated within an initial movement window (300-500 ms after go cue). All black bars are 95% CIs. Performance was generally good across most days, where decoders failed for only a few days. **b** A sequential unimanual control strategy (moving a cursor at a time; solid black line) was compared to simultaneous bimanual control (dashed gray line) over 2 sessions, of which the median target acquisition times are shown. The sequential unimanual control strategy led to faster target acquisitions. **c** RNN decoders were compared to linear decoders on 2 session days. Each point is the median target acquisition time for the corresponding trial type. Solid lines connect points corresponding to the normal bimanual task (consisting of simultaneous dual movements and unimanual single movements). A variation of the task where only unimanual movements were tested (holding the non-active cursor fixed) was used as a control on trial day 1855 to confirm that linear decoders could succeed in a purely unimanual context (dashed lines).

During online control, T5 remarked that sequentially moving the cursors during the bimanual context instead of moving them simultaneously was a more intuitive strategy to employ. To investigate this further, we trained two separate RNNs where one was recalibrated normally as mentioned above, and the other was recalibrated with just unimanual data. On average, the sequential unimanual strategy outperformed the simultaneous bimanual strategy (Fig. 4b, Supplementary Movie 2). Interestingly, the sequential strategy often led to equal performance between unimanual right and unimanual left trials, indicating that the RNN better learned to disentangle the hands when recalibrated on just unimanual movements.

Lastly, we compared linear decoders (LDs) to RNNs for simultaneous two-cursor control. Optimizing linear decoders during online evaluation is difficult since it often requires hand tuning of parameters such as output gain. For the fairest comparison against RNNs, we tested a range of output gain scalars for both LDs and RNNs. However, sweeping the gains did not affect the result of RNNs outperforming the LDs on average (Fig. 4c, Supplementary Movie 3). In fact, the LDs resulted in mostly failed trials due to their inability to isolate control to one cursor (i.e., intended movements of one cursor would inadvertently move the other such that target acquisition was near impossible). As a control, T5 was able to acquire unimanual targets when the non-active cursor was fixed using LDs, indicating that failures during bimanual control were due to the LD’s inability to separate left from right hand control.

### Neural networks leverage laterality information for improved unimanual decoding

Earlier, we found a large neural dimension coding for laterality, which we hypothesized would help identify which hand is moving at any given time – particularly useful if the tuning of the two hands is correlated during unimanual movement. Given that the RNNs outperformed linear decoders during unimanual movement, we sought to dissect the role of laterality information during decoding. First, we compared a simple linear decoder (LD; built via ridge regression) to a simple densely connected feed forward neural network (FFN) to assess each decoder’s ability to use laterality information for unimanual movement decoding. These basic decoders were chosen to mitigate temporal filtering factors (i.e., use of time history as seen with Wiener filters and recurrent networks). That is, which decoder better predicts movement encoded in a single time-bin of neural activity? Using data from unimanual trials, both decoders were trained to convert firing rate input features at a single time-bin (20ms bin) to x- and y-direction velocities for both cursors. Figure 5a shows an example snippet of offline decoded x-direction velocities for unimanual movement of both hands. The FFN outperformed the LD in predicting velocity magnitudes for the left hand, which is consistent with prior results^50^ indicating that ipsilateral representation is generally weaker than contralateral representation (left hand is 48% weaker; see Supplementary Fig. 1d). Figure 5b summarizes offline unimanual decoding performance where the FFN outperformed the LD across all movement dimensions, with the greatest performance boost for unimanual left hand decoding.

**Fig. 5.**
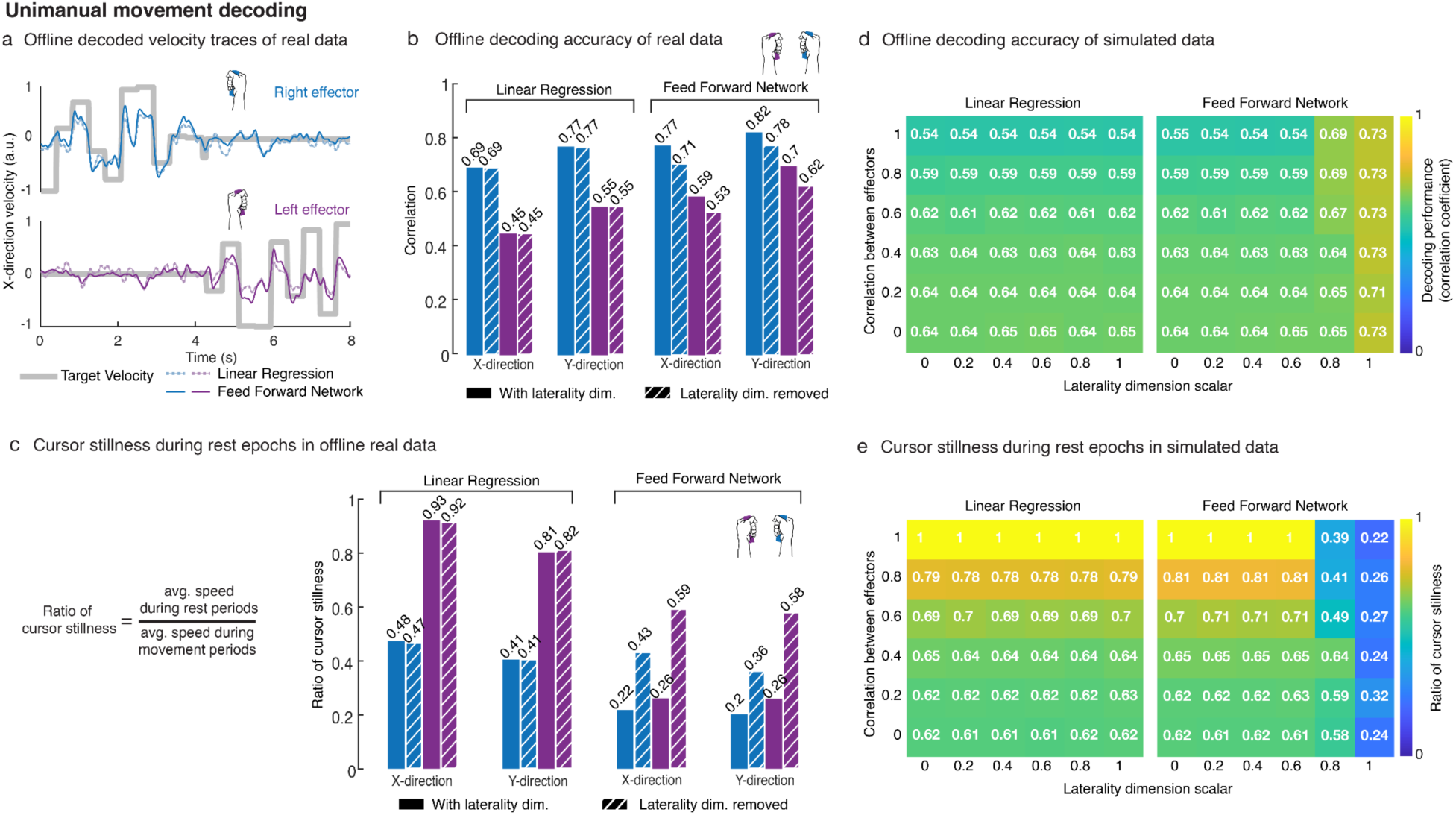
Nonlinear decoders leverage laterality information to disentangle effectors. **a** Offline single-bin decoding on unimanual data. Neural activity was binned (20-ms bins) and truncated to 400 ms movement windows (300-700 ms after go cue). Linear ridge regression (RR) and a densely connected feed forward neural network (FNN; single layer, 512 units) were trained, using 5-fold cross-validation, to decode left and right cursor velocities. Sample 8 s held-out snippets of decoded x-direction velocity traces are shown. **b** Each bar indicates the offline decoding performance (Pearson correlation coefficient) for the RR and FNN decoders across the x- and y-direction velocity dimensions. Striped bars indicate data where the laterality dimension was removed. The FFN outperformed the LD in decoding movements across all dimensions. Removal of the laterality dimension did not affect LD performance and only slightly reduced FFN performance. **c** Cursor stillness is quantified as the ratio of average cursor speed during rest periods to that during movement periods. A rest period is defined as the period in which the other cursor should be active. Lower ratios indicate more cursor stillness while the other cursor is active. The FFN was able to keep each cursor reasonably still, whereas the LD struggled to keep the left cursor still. The laterality dimension was useful to the FFN in keeping the cursors still, however, did not affect the LD. **d** Simulated neural activity during unimanual movement was generated with varying directional tuning correlation between hands and varying laterality dimension size. Each (*i,j*) cell of a matrix indicates the decoding performance (Pearson correlation coefficient) for a synthetic dataset with correlation *i* between hands and a laterality dimension size of *j*. **e** Cursor stillness for the simulated data in panel d is shown. The FFN leveraged the laterality dimension for improved decoding performance and cursor stillness as tuning between the hands became more correlated. The LD was unable to use the laterality information to disentangle the hands.

To further understand the extent to which the decoders used laterality information, we fit and subsequently removed the laterality dimension from neural data (see Methods). Removal of the laterality dimension did not affect decoding performance of the LD whatsoever; however, it did result in a performance hit across all movement dimensions for the FFN (Fig. 5b). Generally, the FFN’s decoding performance was reduced to similar levels to that of the LD’s performance, although the FFN’s left hand decoding was still better than the LD (and its decoded outputs were larger in magnitude; see Supplementary Fig. 3a for distributions of decoded output magnitudes). Additionally, the FFN was better able to isolate movement decoding to the actively moving hand, which we quantified with cursor ‘stillness’ in Figure 5c. On average, the FFN outperformed the LD in keeping the left cursor still during right cursor movement, and vice versa. Removal of the laterality dimension led to a reduction in cursor stillness for the FFN. The LD was unable to keep the left cursor still while the right was active and removal of the laterality dimension did not alter the LD’s ability to keep the cursors still.

To gain deeper insight into the role of laterality information in decoding unimanual movement, we simulated unimanual neural activity with Gaussian noise (see Methods and equations 3,4) where we varied the directional tuning correlation between the hands and varied the size of the laterality dimension. Figure 5d shows decoding performance of LDs and FFNs across the simulated data. As expected, LD performance degraded as the hands became more correlated regardless of the scale of the laterality dimension. Conversely, when the size of the laterality dimension was sufficiently large, the FFNs were able to achieve high decoding performance irrespective of how correlated the hands became. Additionally, we saw that the LDs were unable to use laterality information in keeping the non-active cursor still and cursor stillness degraded as the hands became increasingly correlated (Fig. 5e). The FFNs used laterality information, when it was salient enough, to disentangle the cursors which resulted in increased cursor stillness regardless of how correlated the hands became.

## Discussion

Deep learning algorithms are being increasingly used to improve the performance of real-time BCIs^28, 32, 36, 37, 59, 60^. Prior work investigating deep learning methods for BCIs has reported promising offline results^33, 54–56, 61–64^, although most remain to be evaluated in an online setting due, in part, to the rarity of human BCI data. Here, we tested a deep learning method for real-time BCI control by a person with paralysis. We confronted a challenging nonlinear BCI problem – the simultaneous bimanual control of two cursors – using an RNN, which should be able to exploit the nonlinear structure in the neural data better than linear methods which have been previously used^28, 29, 33, 36^. Consistent with prior work^33, 54–56^, the RNN performed exceedingly well on offline data. However, we found that the high offline performance was due to the RNN overfitting to the temporal structure of the offline data. This, in turn, translated to poor online performance. In response, we altered the temporal structure of the training data which helped the RNN generalize to the online setting, enabling it to far outperform linear methods. Thus, preventing neural networks from overfitting to stereotyped structure in training data may be necessary for translating deep learning methods to real-time BCI control and obtaining the associated performance benefits.

The data alteration method proposed here is one way to approach the problem of neural network overfitting to offline BCI training data, which was accomplished by dilating/compressing smaller snippets of training data and shuffling the order of the modified snippets. There are likely many other methods of helping neural networks generalize to data with less stereotyped structure. For example, there has been a recent compelling approach in NHPs^37^ which recalibrates neural networks by using movement intention estimation techniques motivated by the ReFIT (recalibrated feedback intention-trained) algorithm^39^. In this same study, Willsey et al. deployed a shallow feed-forward network for online BCI control where only 150 ms windows of data were used at each time step. Similar to these short windows of data, we suspect that our data alteration method forced the RNN to learn smaller time histories of data, allowing it to learn the temporal characteristics of bimanual movement-related neural activity without overlearning the specific sequence of behaviors performed during open-loop trials.

An additional useful feature of this method is that it generates synthetic data which helps prevent overfitting to the limited amount of data that is normally collected in human BCI research. Typically, BCI decoder calibration tasks are on the order of minutes and generally do not generate more than a few hundred trials worth of data^12, 13, 20–22, 38, 50, 65^, whereas this method can easily increase this training data quantity by orders of magnitude, which may prevent overfitting (as shown in recent work on handwriting decoding^28^). Future studies could investigate the utility of altering temporal structure in training data across different network architectures and decoding algorithms. Snippet window widths and the quantity of synthetic data are additional hyperparameters that could be further optimized in future work.

In addressing the challenge of decoding bimanual hand movements from neural activity, neural networks were better able to use the nonlinear structure in the neural data compared to linear methods. Laterality information (neural coding for the side of the body) was instrumental in helping the networks distinguish between left and right hand unimanual movements, particularly as neural tuning between the hands became increasingly correlated^50, 66, 67^. Linear decoders cannot leverage laterality information since it is independent of movement direction, resulting here in inadvertent decoded movements of the other effector during unimanual movement.

In this study, we demonstrated bimanual two-cursor control, consistent with a trend towards decoding more challenging behaviors for BCI systems including fine dextrous hand control^37, 42^ and the control of multiple effectors^41, 50, 52^. Deep learning methods will likely be increasingly useful for decoding these complex movements with potentially nonlinear neural representations (as highlighted here for bimanual movements). A key consideration in implementing RNN-based decoders will be to reduce overfitting to stereotyped structure in training data. In sum, altering the temporal and behavioral structure within training data can help translate deep learning methods to real-time BCI control – a potentially necessary step in helping these systems achieve clinical translation.

## Methods

### Study permissions and participant details

This work includes data from a single human participant (identified as T5) who gave informed consent and was enrolled in the BrainGate2 Neural Interface System clinical trial (ClinicialTrials.gov Identifier: NCT00912041, registered June 3, 2009). This pilot clinical trial was approved under an Investigational Device Exemption (IDE) by the US Food and Drug Administrations (Investigational Device Exemption #G090003). Permission was also granted by the Stanford University Institutional Review Board (protocol #20804) and the Mass General Brigham IRB (protocol #2009P000505).

Participant T5 is a right-handed male (69 years of age at the time of study) with tetraplegia due to cervical spinal cord injury (classified as C4 AIS-C) which occurred approximately 9 years prior to enrollment in the clinical trial. In August 2016, participant T5 had two 96-channel intracortical microelectrode arrays (Blackrock Microsystems, Salt Lake City, UT; 1.5 mm electrode length) placed in the hand knob area of the left (dominant) precentral gyrus. The hand knob area was identified by pre-operative magnetic resonance imaging (MRI). Supplementary Figure 1a shows array placement locations registered to MRI-derived brain anatomy. T5 has full movement of the face and head and the ability to shrug his shoulders. Below the level of spinal cord injury, T5 has very limited voluntary motion of the arms and legs. Any intentional movement of the body below the level of injury is referred to as being “attempted” movement where small amplitude movements were intermittently observed.

### Neural data processing

Neural signals were recorded from two 96-channel Utah microelectrode arrays using the NeuroPort^TM^ system from Blackrock Microsystems (see [^12^] for basic setup). First, neural signals were analog filtered from 0.3 to 7.5 kHz and subsequently digitized at 30kHz with 250 nV resolution. Next, common mode noise reduction was accomplished via a common average reference filter which subtracted the average signal across the array from every electrode. Finally, a digital high-pass filter at 250 Hz was applied to each electrode prior to spike detection.

Spike threshold crossing detection was implemented using a -3.5 x RMS threshold applied to each electrode, where RMS is the electrode-specific root mean square of the time series voltage recorded on that electrode. Consistent with other recent work, all analyses and decoding were performed on multiunit spiking activity without spike sorting for single neuron activity^68–70^.

### Session structure and two-cursor tasks

Neural data was recorded from participant T5 in 3-5 hour “sessions”, with breaks, on scheduled days (see Supplementary Table 2 for a comprehensive list of data collection sessions). T5 either sat upright in a wheelchair that supported his back and legs or laid down on a bed with his upper body inclined and head resting on a pillow. A computer monitor was placed in front of T5 which displayed two large circles indicating targets (one colored purple and one colored white) and two smaller circles indicating cursors with corresponding colors. The left cursor was labeled ‘L’ and colored purple and the right cursor was labeled ‘R’ and colored white.

During the open-loop task, the cursors moved autonomously to their designated targets in a delayed-movement paradigm. On each trial, one of three movement types were cued randomly: (1) bimanual (simultaneous movement of both cursors), (2) unimanual right (only right cursor movement), and (3) unimanual left (only left cursor movement). Each trial began with a random delay period ranging from 1-2 seconds where lines appeared and connected each cursor to its intended target. During the delay period, T5 would prepare the movement. After the delay period, indicated by a beep sound denoting the ‘go’ cue, the lines disappeared and the cursors moved to their targets over a period ranging 1-2 seconds in length, where cursor movement was governed by a minimum-jerk trajectory^50, 71^ (black velocity profile in Fig. 3b). Both cursors arrived at their intended target at the same time. T5’s attempted movement strategy was to imagine that his hands were gripping joysticks (as illustrated in Fig. 1a) and to push on each joystick to control the corresponding cursor’s motion. The end of each trial was indicated by another beep sound where T5 was instructed to stop all attempted movements and to begin preparing for the next trial’s movement.

The closed-loop tasks generally mimicked the open-loop task except that the cursors were controlled via neural decoders (either an RNN or linear decoder) instead of having prescribed motion to their targets. During each closed-loop trial, T5 had a maximum of 10 seconds to acquire both targets. Target acquisition was defined as both cursors simultaneously dwelling within their intended target for an uninterrupted duration of 500 ms. If any one cursor moved outside of its target before the dwell period elapsed then the dwell timer was restarted. Both targets were illuminated blue during a proper simultaneous dwell (see Supplementary Movie 1). If the targets were not successfully acquired within the 10 second timeout period then the trial was considered failed.

An “assisted” version of the closed-loop task was often used for decoder recalibration prior to true closed-loop evaluation blocks. Assistance was provided in the form of “error assistance” and/or “push assistance”. Error assistance^11, 72^ was accomplished by attenuating velocity commands in the dimensions orthogonal to each cursor’s straight-line path to the respective target. The attenuation factor was determined by a scalar value ranging from 0-1 where 0 provided no error assistance and 1 would remove all orthogonal velocity commands resulting in cursor movement along the line to the target. Push assistance was given for each cursor via adding a unit velocity vector in the direction of the corresponding target (referred to as “push vector”) which was scaled by the decoded cursor speed (magnitude of the velocity vector). The degree of push assistance was also governed by a scalar value ranging from 0-1 where 0 provided no push assistance and 1 would scale the push vector to the size of the decoded cursor velocity vector. The point of push assistance was to reinforce movement to the intended target by only aiding when the participant was trying to move. The amount of push and error assistance on each block was governed by the experimenter to ensure that the participant was able to acquire most, if not all, targets for recalibration purposes.

Since performance during recalibration was generally suboptimal, unimanual trials would often result in movement of both cursors which then would require bimanual control to correct cursor deviation. This was not ideal when considering the balance of training data for trial and movement type. To address this, we instituted a “lock mode” where the non-active cursor’s motion was fixed so that the participant was able to focus on the cursor which was cued to move during unimanual trials.

### Offline population-level analyses

#### Cross-validated estimates of neural tuning strength and tuning correlation between effectors

We used cross-validated estimates of Euclidean distance for the quantification of neural tuning strength and other statistics requiring Euclidean distance, such as Pearson’s correlation between groups of linear model tuning coefficients. These methods are discussed in greater detail in our prior report^50^ (Willett*, Deo*, et al. 2020; see code repository https://github.com/fwillett/cvVectorStats).

Tuning strength was quantified using a cross-validated implementation of ordinary least squares regression (cvOLS.m) to estimate the magnitude of columns of linear model coefficients. Tuning coefficients were found using the following model:

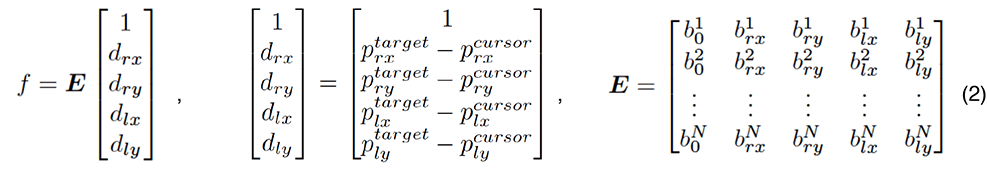

Here, *f* is the N x 1 firing rate vector for a single time step where N is the number of electrode channels. *E* is an N x 5 matrix of mean firing rates (first column; superscript denotes electrode number) and directional tuning coefficients (second to fifth columns; superscript is electrode number and subscript represents the hand as *r* or *l* and movement as the *x*- or *y*-direction). Variables *d_rx_*, *d_ry_*, *d_lx_* and *d_ly_* of the predictor vector represent the *x* and *y* components of the right (*r*) and left (*l*) hand’s intended movement defined as the corresponding difference between target position (*p* terms with superscript ‘*target’*) and cursor position (*p* terms with superscript *‘cursor’*). *E* was fit via 5-fold cross-validated ordinary least-squares regression using 20-ms binned data within a window from 300 to 700 ms after the go cue across all trials. This was accomplished by “stacking” the response (firing rate) and predictor vectors horizontally across all candidate timesteps. Block-wise means were calculated and subtracted from all neural data prior to analyses to adjust for nonstationarities and neural drift over time^73, 74^.

The data used in Figure 2 were from 5 session days (trial days 1776, 1778, 1792, 1881 and 1883) where we were able to collect large amounts of unimanual and bimanual open-loop data (since cross-validation requires each fold to have enough data to properly estimate regression coefficients). For each session day, we grouped consecutive blocks together in pairs to reach around 40 repetitions, at least, per trial type (unimanual right, unimanual left, and bimanual). Within each block set, we used the cvOLS function to compute the coefficient vectors and their magnitudes for each movement type. That is, we fit a separate model to all unimanual right trials, all unimanual left trials, and all bimanual trials. Notice that fitting the unimanual models reduces the encoding matrix *E* to three columns (e.g., the last two columns related to the left hand are removed when fitting for unimanual right movement). We defined tuning strength for each hand (right or left) under the unimanual or bimanual contexts by averaging over the corresponding model’s x- and y-direction coefficient vector magnitudes. Ratios of these tuning strengths between models across each pair of block sets are reported in Figure 2b (gray dots; sample size of 25). Tuning correlation in Figure 2c was quantified by computing the (cross-validated) Pearson correlation between corresponding x or y- direction coefficient vectors between models (gray dots indicate correlations between models, as listed on the x-axis, across all block-sets). Correlations were computed using the cvOLS function. The x- and y- direction correlations are shown separately since the hands are more correlated in the y-direction and anti-correlated in the x-direction, as we have previously shown^50^, which would result in nullifying effects if correlations were averaged across direction dimensions.

#### Principal component analysis (PCA) of laterality coding

We used PCA to visualize the neural activity in a lower-dimensional space as illustrated in Figure 2d. Using data from one of the sessions (trial day 1881) described above (20-ms binned, block-wise mean removed, Z-scored), we computed each trial’s average firing rate vector within the 300-700 ms window after the go cue. We then stacked each trial’s N x 1 firing rate vector horizontally resulting in an N x T matrix where T is the number of trials. PCA was performed on this monolithic matrix and each firing rate vector was subsequently projected onto the top two principal components (PCs) as illustrated in the left panel of Figure 2d. The single-trial projections were colored by the trial type (unimanual right trial, unimanual left trial, or bimanual trial) to show how the data clustered. Next, we projected each trial’s binned firing rates across time (-500 ms to 1.5 s relative to the go cue) onto the top PC to visualize a population-level peristimulus time histogram. Each thin line corresponds to a single trial’s projection, colored by trial type, and the bold lines are the mean projections shaded with 95% confidence intervals computed via bootstrap resampling.

In order to quantify the size of laterality-related tuning, we used a variation of demixed principal component analysis^57^ (dPCA; Kobak et al., 2016; https://github.com/machenslab/dPCA). A central concept of dPCA is marginalizing the neural data across different sets of experimentally manipulated variables, or factors. Each marginalization is constructed by averaging across all variables that are not in the marginalized set, resulting in a data tensor that captures the effect of the factors on the neural activity. dPCA then finds neural dimensions that explain variance in each marginalization alone, resulting in a useful interpretation of neural activity according to the factors. Leveraging the existing dPCA library, we implemented a cross-validated variance computation to reduce bias by splitting the data into two sets, marginalizing each set, element-wise multiplying the marginalized matrices together, and summing across all entries. The data was marginalized over the following four factors: laterality, movement direction, laterality x movement direction interaction, and time. For each dataset used in Figure 2b,c (trial days 1776, 1778, 1792, 1881 and 1883), we computed the cross-validated variance in the aforementioned factors. The bar plots in Figure 2d (rightmost panel) summarize the average cross-validated marginalized variance for each factor (labeled along the x-axis) across all 5 sessions (gray dots).

### Single electrode channel tuning

To assess neural tuning to unimanual or bimanual movement on a given electrode as seen in Supplementary Fig. 1, we used a 1-way ANOVA on firing rates observed during directional hand movements within each movement context. This analysis was performed on the same dataset used in Figure 1 (trial day 1750). We first computed the average firing rate vector for each trial within the 300 to 700 ms window relative to the go cue. Next, we separated each of the computed average firing rate vectors into the following sets: unimanual right trials, unimanual left trials, and bimanual trials. Within each set, we grouped the vectors into their respective movement direction (4 directions defined by each quadrant in the unit circle) for each hand. Grouping the bimanual trials for right hand movement direction ignored left hand movement direction and vice versa. This resulted in 4 total sets of firing rate vectors grouped by their respective hand’s movement direction (unimanual right directions, unimanual left directions, bimanual right directions, and bimanual left directions) and a separate 1-way ANOVA was performed within each set. If the p-value was less than 0.00001, the electrode was considered to be strongly tuned to that movement context (unimanual or bimanual). To assess the tuning strength of each strongly tuned electrode, we computed FVAF (fraction of variance accounted for) scores^50, 65^. The FVAF score was computed using the following equations:

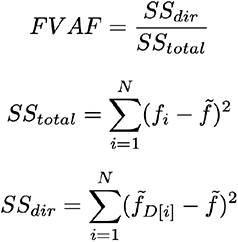

Here, *SS_total_* is the total variance (sum of squares), *SS_dir_* is the movement direction-related variance, *N* is the total number of trials, *f_i_* is the average firing rate vector for trial *i*, *f̃* is the average firing rate vector across all trials within the set, and *f̃_D_*_[*i*]_is the average firing rate vector for the particular movement direction cued on trial *i*. FVAF scores range from 0 (no direction-related variance) to 1 (all variance is direction-related).

### Training data augmentation via dilation and randomization of training snippets

The raw data as formatted for RNN training took the form of an input ‘feature’ data tensor of shape S x T x N and an output ‘target’ tensor of shape S x T x R. Here, S is the number of training snippets, T is the number of time points in a snippet, N is the number of electrode channels, and R is the number of response or output variables. The input tensor consisted of neural data which was binned at 20 ms, block-wise mean removed, and Z-scored. The output tensor contained the cursors’ velocities and movement context signals which were also binned at 20 ms (see Fig. 3 and Supplementary Fig. 2). Typically, we held our training snippet length at 10 s (T=500 at 20-ms bins). We generated a large number of synthetic training snippets by splicing together smaller pieces of the data stream which were also dilated in time and random in order.

Our objective was to generate an augmented dataset which was balanced across movement direction and movement type. We defined 4 gross movement directions corresponding to each quadrant of the unit circle and movement type was defined as unimanual, bimanual, or no-movement. The types of no-movement were further subdivided into the following groups: (1) unimanual right delay period, (2) unimanual left delay period, (3) bimanual delay period, and (3) rest. This distinction in types of no-movement was so that we may equally account and balance for preparatory activity as well as rest activity. The training data was preprocessed to label each data sample’s movement quadrant per hand and movement type. We generated roughly 2000 synthetic training snippets (each snippet of 10 s length) for training, which was chosen based on the time it took to perform the augmentation during an average experiment session (10-15 minutes). A synthetic 10 s training snippet was generated by appending dilated/compressed clips of raw data. Each raw data clip was selected to begin at a random time point, varied in duration (ranging between 0.2 to 0.8 times the 10s total snippet length), and had an associated dilation/compression factor *d_f_* drawn from a uniform distribution over the interval [0.5, 2], where *d_f_* = 1 indicates no change, *d_f_* < 1 indicates compression, and *d_f_* >1 indicates dilation. For a candidate clip to be considered valid, it had to abide by the current balancing record which was kept across all of the aforementioned movement conditions. Generally, the input data was balanced to achieve a sufficient amount of data for each of the movement types. If the candidate clip did not meet the balancing requirements, then another random clip was drawn. Linear interpolation was used to either compress or stretch both the input and output clips of raw data based on the dilation/compression factor (e.g., a *d_f_* of 0.5 would compress a clip array of length 60 into an array of length 30 by sampling every other element of the original clip). The data augmentation method generated both a training and held-out validation set that did not contain overlapping data. The input data was split into a training and validation set in advance, then from these isolated pools augmented sets of training data could be created.

### Online recurrent neural network decoding of two-cursors

We used a single-layer gated recurrent unit (GRU) recurrent neural network architecture to convert sequences of threshold crossing neural firing rate vectors (which were binned at 20 ms and Z-scored) into sequences of continuous cursor velocities and discrete movement context signals. The discrete context signals coded for which movement (unimanual right, unimanual left, bimanual, or no movement) occurred at that moment in time and enabled the corresponding cursor velocity commands to be gated. We used a day-specific affine transform to account for inter-day changes in neural tuning when training data were combined across multiple days. The RNN model and training was implemented in *TensorFlow v1*. The online RNN decoder was deployed on our real-time system by extracting the network weights and implementing the inference step in custom software. The RNN inference step was 20 ms. A diagram of the RNN is given in Supplementary Fig. 2.

Before the first day of real-time evaluation, we collected pilot offline data across 2 session days (trial days 1752 and 1771) comprising 1 hour of 780 total trials (balanced for unimanual and bimanual trials) which were combined to train the RNN. All training data were augmented to generate around 2000 training snippets of ten second length amounting to roughly 6 hours of data (balancing equally for each movement type). We tuned the initial RNN model’s hyperparameters (input noise, input mean drift, learning rate, batch size, number of training batches, and L2-norm weight regularization) via a random search deployed across 100 RNNs. On each subsequent day of real-time testing, additional open-loop training data were collected (approximately 25 minutes of 280 trials; roughly 6 hours of 30K trials after augmentation) to recalibrate the RNN which was subsequently used to collect 4 assisted closed-loop blocks (5 minutes each) for a final recalibration. For each RNN recalibration, all data that were used for training up until that point in time were included, where 40% of training examples were from the most recently collected dataset and the remaining 60% of training examples were evenly distributed over all other previously collected datasets. During recalibration periods in which the RNN was training, firing rate means and standard deviations were updated via an elongated open-loop block (8-minutes in length) which were used to Z-score the input firing rates prior to decoding. This RNN training protocol was used for the unimanual and simultaneous bimanual data presented in Figure 4a. In total, performance was evaluated across 6 days (trial days 1752, 1771, 1776, 1778, 1790, 1792) with each day containing between 4-8 blocks (5 minutes each) with balanced trials across each movement context.

The RNN training varied slightly for the ‘sequential bimanual’ data presented in Figure 4b. The base RNN (prior to the first day of real-time evaluation) was calibrated in the same fashion as mentioned above, however each subsequent dataset used for recalibration consisted of just unimanual trials and no bimanual trials. Data from two evaluation sessions (trial days 1881 and 1883) were used for Figure 4b.

The data augmentation panels of Figure 3d,e were generated based on data from two session days (trial days 1867 and 1869). The two separate RNNs used were trained only on the data gathered during those sessions and did not include any historical data to focus on the effects of our data augmentation technique. One RNN was trained with data that was augmented and the other RNN was trained on the raw non-augmented data. The open-loop results and sample speed traces shown in Figure 3d,e are from trial day 1869.

#### Online two-cursor control performance assessment

Online performance was characterized by time-to-acquisition and angular error. Time-to-acquisition for a trial was defined as the amount of time after the go cue in which the targets were successfully acquired. Angular error was defined as the average difference between movement direction within the 300 to 500 ms window after the go cue to capture the ballistic portion of each movement prior to any error correction. Each trial timed out at 10 seconds, after which the trial was considered failed.

#### Comparing linear regression and RNN decoding

We tested a range of output gains for the comparison of online linear decoders and RNNs used for Figure 4c (includes data from trial days 1853 and 1855) to ensure that performance differences were not due to variation in decoded output magnitudes. The range of gain values was determined on each session day by a closed-loop block (preceding data collection) where the experimenter hand-tuned values until the participant’s control degraded. Hand-tuning of gain values was done for the linear decoder and RNN, separately. Each session day had 4-5 equally spaced gain values for each decoder. For the data presented in Figure 4c, we averaged over all swept gains to summarize performance for each decoder since it turned out that the result was not affected by what gain was used (e.g., linear decoder results include data from each swept gain).

### Offline single-bin decoding of real and simulated unimanual data

#### Real and simulated neural data for unimanual movement

The real unimanual dataset analyzed for Figure 5a,b,c was from trial day 1883. The data were binned (20- ms bins), block-wise mean removed, and each trial truncated to 400 ms movement windows (300 to 700 ms after the go cue). In keeping with standard BCI decoding practice and to focus on directional movement decoding, we defined the velocity target for each time step as the unit vector pointing from the cursor to the target, resulting in discrete velocity steps as seen in Figure 5a (thick gray lines).

When generating synthetic data for simulations, we attempted to match the ‘functional’ signal-to-noise ratio (fSNR) of the real dataset for a more practical comparison. The fSNR decomposes decoder output into a signal component (a vector pointing at the target) and a noise component (random trial-to-trial variability). We first generated the decoder output using a cross-validated linear filter to predict a point-at-target unit vector *y_t_* (normalized target position minus cursor position) given neural activity as input.

We then fit the following linear model to describe the decoder output:

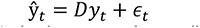

Here, *y_t_* is the 2 x 1 point-at-target vector, *ŷ_t_* is the cursor’s predicted velocity vector at timestep *t*, *D* is the 2 x 2 decoder matrix, and ϵ*_t_* is the 2 x 1 vector of gaussian noise at timestep t.

We computed the functional SNR (*fSNR*) as:

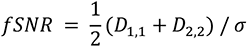

Here, *D*_1,1_ and *D*_2,2_ are the diagonal terms (subscripts refer to row i and column j) of the 2 x 2 *D* matrix, and σ is the standard deviation of *ɛ* (averaged across both dimensions). We estimated D by least squares regression. We estimated σ by taking the sample standard deviation of the model error. Intuitively, the numerator describes the size of the point-at-target component of the decoder output, and the denominator describes the size of the trial-to-trial variability.

To simulate neural activity, we used the laterality encoding model in equation 4 where we varied the directional tuning correlation between the hands and the size of the laterality dimension (as labeled along the x- and y-axes of Fig. 5d,e). We began by generating a synthetic target dataset containing unimanual velocities for the left and right hands. The synthetic targets consisted of approximately 2000 unimanual right trials and 2000 unimanual left trials. Trial lengths were 400 ms in duration to match the real dataset and binned in 20-ms bins. The synthetic target data were balanced across 8 movement direction wedges evenly distributed throughout the unit circle (see x-axes in Fig. 1c for direction wedges). Specifically, a uniformly random unit velocity vector was generated within a direction wedge for each trial ensuring even distribution across all wedges for both hands. Essentially, the synthetic targets resembled the sample real-data targets seen in Figure 5a (thick gray lines). Next, we generated random tuning coefficients (*b* terms in eq. 4) for 192 synthetic neurons by sampling from a standard normal distribution. The population-level tuning vectors were then scaled to match the magnitudes of corresponding tuning vectors from the real dataset (using cvOLS). We then enforced a correlation (which was swept, see y-axes of Fig. 5d,e) between the x-direction tuning vectors for both hands as well as the y-direction vectors. Next, we passed the synthetic velocity targets through the tuning model to compute the population-level firing rates for each time bin. The fSNR for each hand was matched to the real data via adding gaussian noise to each individual channel (sweeping the standard deviation parameter) until the fSNRs of the synthetic data was close to that of real data. The simple noise model is described as follows:

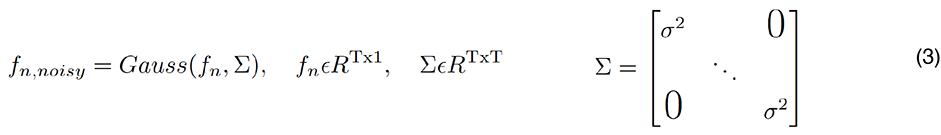

Here, *f_n_* is a T x 1 time-series vector of firing rates for channel *n* where T represents the number of 20-ms time bins, Σ is the T x T diagonal covariance matrix, and σ is the standard deviation. This was a simple noise model with a diagonal covariance matrix used for all channels (i.e., the same σ was used for all channels) . We understand that more sophisticated noise models could have been used, but our simplified approach was well enough suited for single-bin decoding where one can assume independence between time bins which is further explained in Supplementary Figure 4. After matching the fSNRs, we scaled the laterality coefficient vector where a value of 0 removed the laterality dimension completely, and a value of 1 matched the laterality coefficient magnitude of the real data. Finally, we enforced that no firing rates were below zero by clipping negative firing rates to 0.

#### Linear ridge regression and feed forward neural network for single-bin decoding

The real data was split into 5-folds for cross-validation with balanced unimanual right and unimanual left time steps of data within each fold. Cross-validation was necessary for the real dataset since the number of trials was relatively small (482 total trials) in comparison to the simulated dataset (4000 total trials).The simulated datasets were large enough and balanced in terms of trial types that in addition to cross-validation during decoder training, performance was based on completely held out test sets (20% of total simulated data) which were also balanced for trial type.

Simple linear ridge regression was performed on the real and simulated datasets using a neural decoding python package (https://github.com/KordingLab/Neural_Decoding) and the Scikit-Learn library (RidgeCV function). The ridge parameter was swept until decoding performance (measured as the Pearson correlation coefficient) was maximized across all output dimensions. Each feed forward neural network (FFN) was designed as a single densely connected layer of 512 units (*TensorFlow v.1*). The FFNs were initialized with random weights and model parameters were tuned based on an offline hyperparameter sweep on pilot data. All decoders were trained to convert firing rate input features (N x 1 vector) at a single time-bin (20ms bin) to x- and y-direction velocities for both cursors (4 x 1 velocity vector at each time step).

#### Removing laterality information from real unimanual data

Laterality information was removed from real unimanual data by first fitting the linear tuning model below using cross-validation:

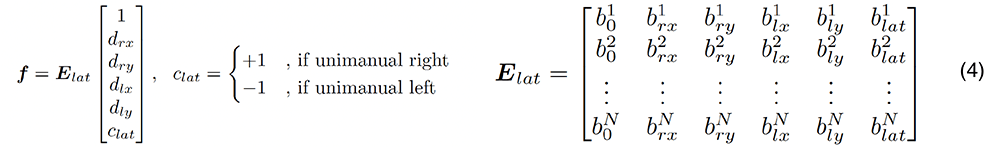

Here, the model resembles that in equation 2 except with the addition of a laterality predictor variable (*c_lat_*) which is +1 for unimanual right movement or -1 for unimanual left. There is an additional column of coefficients (*b_lat_* terms) in the encoding matrix *E*. After this model was fit, the neural activity was projected onto the laterality dimension (last column vector of *E*) and the projected neural activity was subsequently subtracted from the original neural activity. To ensure that laterality information was sufficiently removed, we built another linear filter on the laterality-removed data and confirmed that the laterality coefficients were all zero.

## Acknowledgements

We thank participant T5 and his caregivers for their generously volunteered time and dedicated contributions to this research as part of the BrainGate2 pilot clinical trial, Sandrin Kosasih, Beverly Davis, and Kathy Tsou for administrative support, Erika Woodrum for the drawings in Figs. 1a, 3a, and Elias Stein for help in coding the data augmentation. Support provided by the NIH National Institute of Neurological Disorders and Stroke (U01-NS123101); NIH National Institute on Deafness and Other Communication Disorders (R01-DC014034); Wu Tsai Neurosciences Institute; Howard Hughes Medical Institute; Larry and Pamela Garlick; Office of Research and Development, Rehabilitation R&D Service, US Department of Veterans Affairs (A2295R, N2864C).

## Author contributions

We have included a graphical representation of author contributions as a heatmap below:

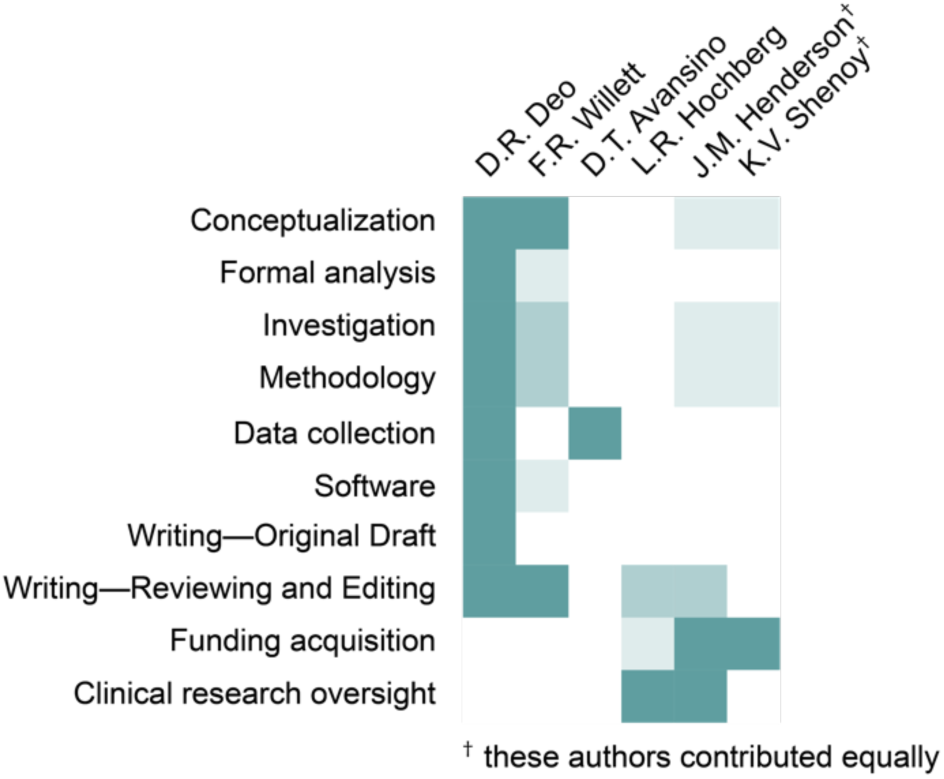

**Supplemental Movie 1 | Simultaneous bimanual control of two cursors via RNN decoding.** In this movie, participant T5 uses a BCI to control two cursors in real-time to targets on a computer monitor. An RNN converts neural activity into velocities for both cursors at each timestep. On each trial, one of three movement types are cued randomly: (1) bimanual (simultaneous movement of both cursors), (2) unimanual right (only right cursor movement), or (3) unimanual left (only left cursor movement). Each trial begins with a ‘prepare’ segment (of random duration) where lines connect each cursor to its intended target. T5 prepares to move during this segment but does not attempt movement until the lines disappear, indicating the ‘go’ cue. Successful target acquisition occurs when both cursors simultaneously dwell within their designated target (illuminates blue) for an uninterrupted period of 0.5 s. A trial times out at a maximum of 10 s. The RNN decoder is enabled at all times. This experiment block was recorded during a performance evaluation session reported in Figure 4 (trial day 1778).

**Supplemental Movie 2 | Sequential unimanual movement vs. simultaneous bimanual movement.** The same as Supplemental Movie 1, except T5 uses two different movement strategies: (1) sequential unimanual (moving one cursor at a time), and (2) simultaneous bimanual (moving both cursors simultaneously). A separate RNN decoder is used for each movement strategy. The RNN used for the simultaneous bimanual strategy is trained normally (just like in supplemental video 1) with both unimanual and bimanual data. The RNN used for the sequential unimanual strategy is trained only with unimanual trials. Both experiment blocks were recorded during a performance evaluation session reported in Figure 4b (trial day 1883).

**Supplemental Movie 3 | RNN vs. linear decoder for two-cursor control.** The same as Supplemental Movie 1, except with only unimanual trials. An RNN decoder is compared to a linear decoder for online control of two cursors. This task was limited to unimanual trials to focus on the differences between decoders. Both experiment blocks were recorded during a performance evaluation session reported in Figure 4c (trial day 1855).

**Supplemental Movie 4 | Online two-cursor control with raw and temporally altered training data.** Same as Supplemental Movie 1, except with only unimanual trials. During this task, one cursor is cued on any given trial where the other cursor stays ‘locked’ in place. This version of the task was used to focus on the differences between decoders. One decoder was trained with raw training data and the other decoder was trained with temporally altered training data. Both experiment blocks were recorded during a performance evaluation session reported in Figure 3e (trial day 1869).

---

**Supplementary Fig. 1.**
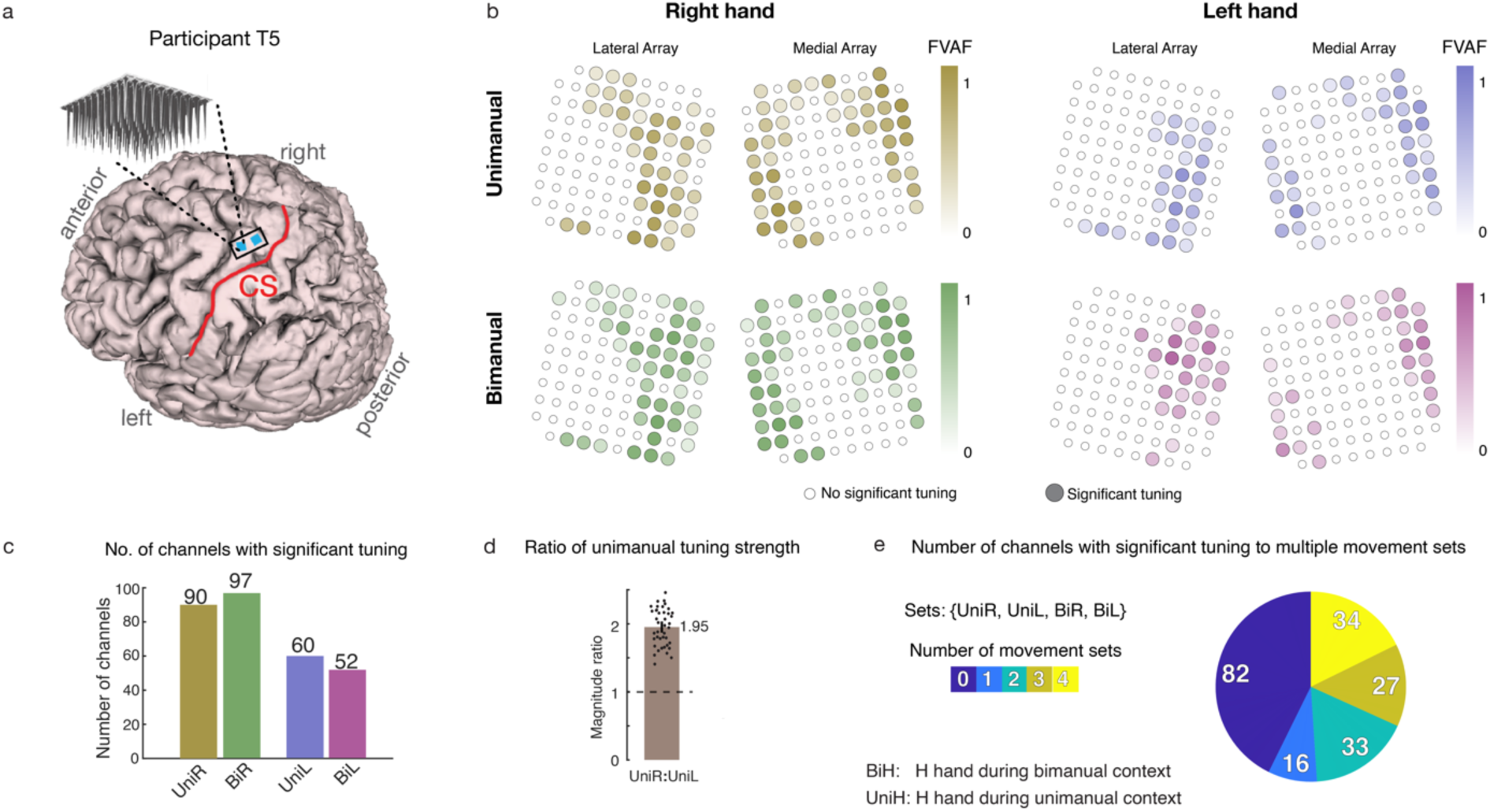
Tuning to unimanual and bimanual movement is intermixed within electrodes and has no clear somatotopic pattern. **a** Participant T5’s MRI-derived brain anatomy and microelectrode array locations. Microelectrode array locations were determined by co-registration of post-operative computed tomography (CT) images with preoperative MRI images. **b** The strength of each electrodes’ tuning to right or left hand movement during unimanual and bimanual movement contexts is indicated with a shaded color (darker colors indicate more tuning). Tuning strength was quantified as the fraction of total firing rate variance accounted for by changes in firing rate due to the movement conditions (unimanual/bimanual). Small white circles indicate electrodes that had no significant tuning to that movement context as governed by a 1-way ANOVA. Broad spatial tuning to all movement categories can be seen across all arrays. **c** Bar plots indicate the number of electrodes that were significantly tuned to each movement context as computed in (a). Results show greater preference for right hand tuning across both movement contexts. **d** Ratio of unimanual tuning strength between the right and left hand. Tuning strength was computed using an unbiased estimate of neural distance between tuning coefficient vectors. The right hand had almost twice as strong tuning than the left hand. **e** Pie chart summarizes the number of electrodes that had statistically significant tuning to each possible number of movement sets (from 0 to 4).

**Supplementary Fig. 2.**
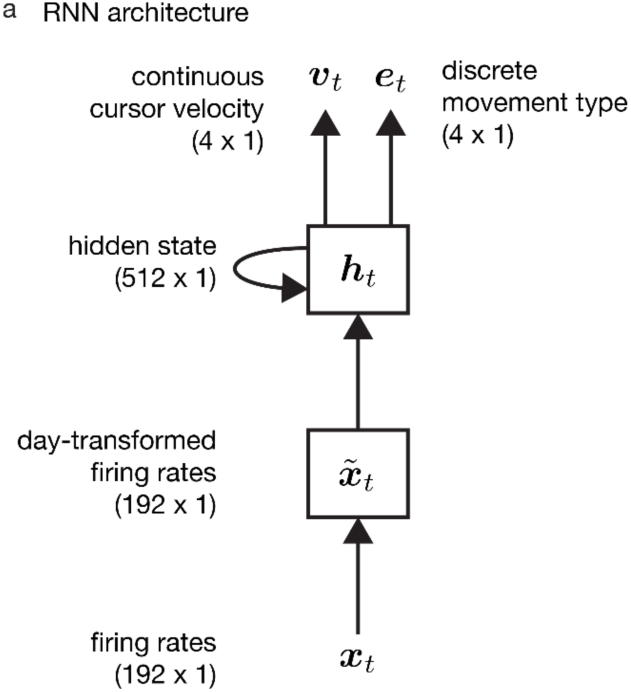
Diagram of the RNN architecture. **a** We used a single-layer RNN with 512 gated recurrent units (GRUs; *h_t_*) to transform neural firing rates (*x_t_*) binned in 20 ms to continuous cursor velocities (*ν_t_*) and discrete movement signals (*e_t_*). The *v_t_* vector describes the x- and y-direction velocities for the right (first two dimensions) and left (last two dimensions) cursors at that moment in time (*t*), and *e_t_* is a one-hot vector (only one dimension is high at any given time) which codes for the type of movement that the RNN detects (unimanual right, unimanual left, bimanual, or no movement) at that time point. Note that we used a day-specific affine transform on the input firing rate vector *x_t_* to account for day-to-day changes in neural activity.

**Supplementary Fig. 3.**
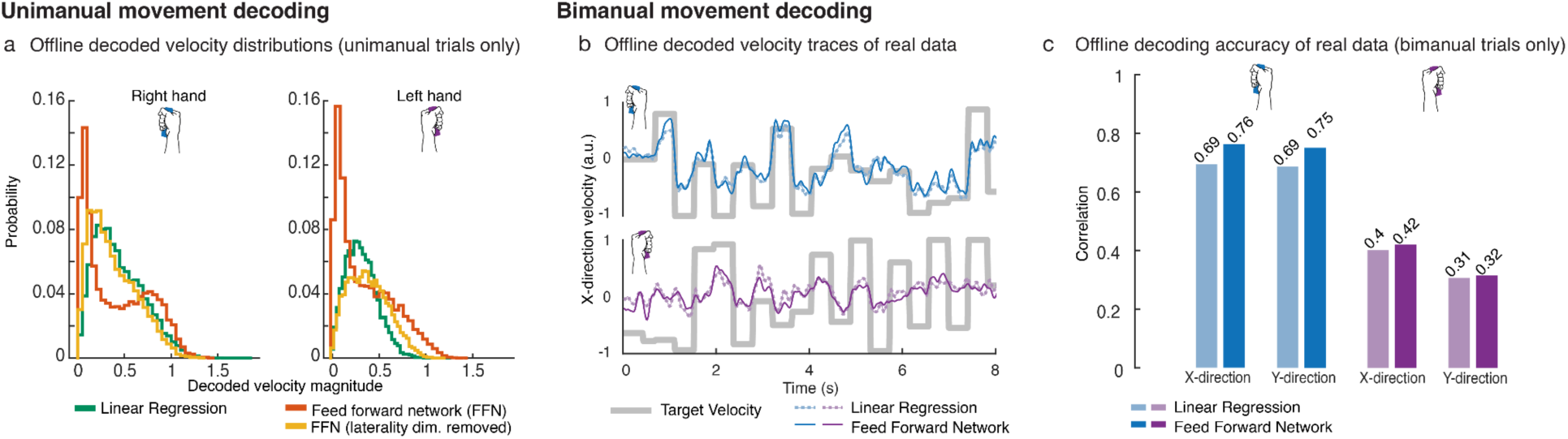
Offline unimanual and bimanual decoding. **a** Distributions of decoded velocity magnitudes during unimanual movement (related to Fig. 5a,b). The feed forward neural network (FFN) was able to decode higher velocity magnitudes than the linear decoder. Removal of the laterality dimension resulted in less decoded velocities near 0 for the FFN, indicating worse cursor stillness without laterality information. **b** Offline single-bin decoding on bimanual data. Neural activity was binned (20-ms bins) and truncated to 400 ms movement windows (300-700 ms after go cue). Linear ridge regression (RR) and a densely connected FFN (single layer, 512 units) were trained, using 5-fold cross-validation, to decode left and right cursor velocities. Sample 8 s snippets of decoded x-direction velocity traces are shown. **c** Each bar indicates the offline decoding performance (Pearson correlation coefficient) for the RR and FNN decoders across the x- and y-direction velocity dimensions. Generally, right hand decoding accuracy was higher than left hand decoding accuracy during bimanual movement.

**Supplementary Fig. 4.**
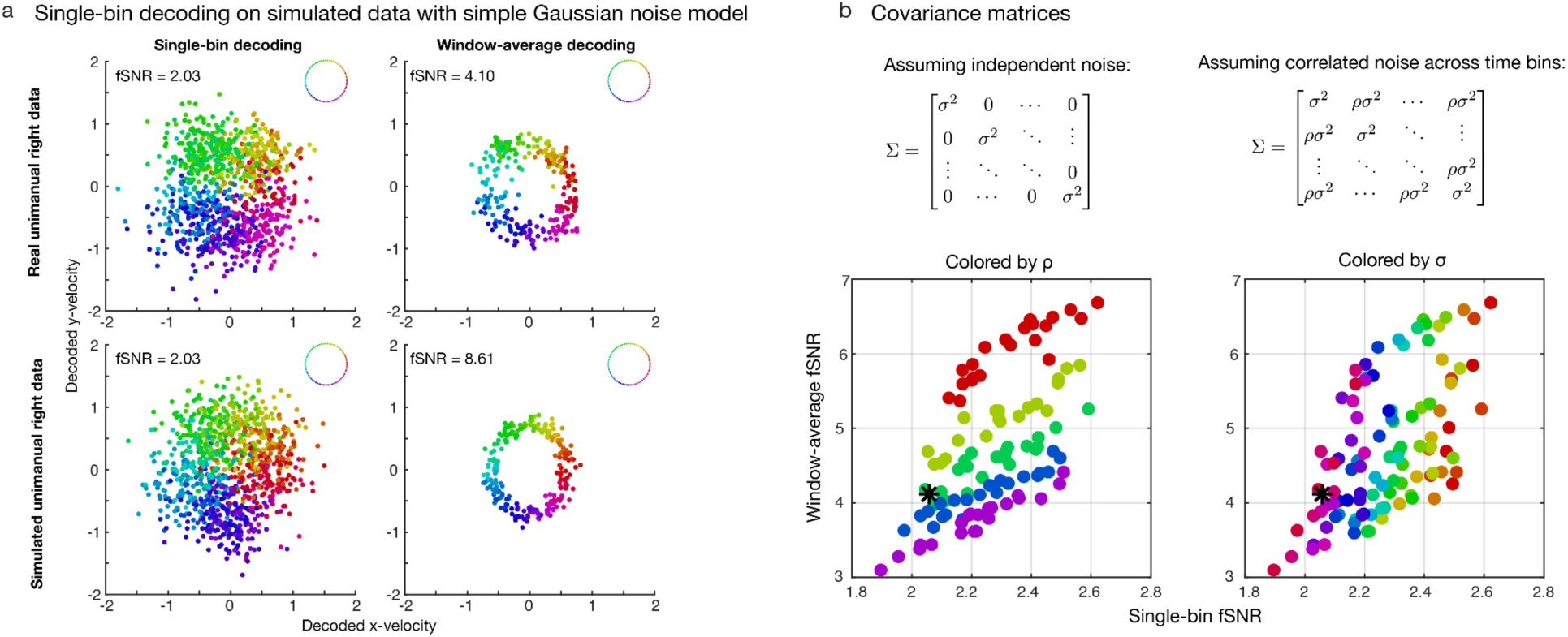
Decoding simulated data with a simple Gaussian noise model. **a** Using a functional signal-to-noise ratio (fSNR; see Methods) metric, we quantify single-bin and window-average decoding performance on real data (top row) and simulated data (bottom row). The simulated data was created using a simple Gaussian noise model (using the left covariance matrix in b) which assumes independence between firing rates across time bins for any given electrode channel. Decoders were built using cross-validated linear regression on 20-ms binned data in the movement window from 300 to 700 ms after the go cue of each trial. The single-bin decoders were calibrated on each 20-ms bin of data, whereas the window-average decoders were calibrated on the averaged activity within the 400-ms window. Each dot represents the decoded x- and y-direction velocity in either each 20-ms time bin (left column) or each 400-ms window of a trial (right column). The color of each dot corresponds to the true target direction of movement indicated by the keys in the upper right of each panel. In this example, the simulated data was generated to match the single-bin fSNR of real unimanual right hand movement data (2.03). Notice that although the single-bin fSNRs of both the real and simulated datasets match, the window-average fSNRs differ quite significantly. **b** The simple gaussian model on the left assumes independent noise, whereas the covariance matrix on the right assumes correlated noise across time bins. σ is the standard deviation, and ρ is the correlation coefficient. **c** In order to match the window-average fSNR, one could use the correlated noise model and sweep the covariance parameters until a window-average fSNR is met. The scatter plots indicate the single-bin and window-average fSNRs of synthetic datasets created by sweeping a range of both σ and ρ parameters in the Gaussian model with correlated noise. Black stars indicate the real data’s single-bin and window-average fSNR as seen in panel a. Both plots are identical except for the way in which the points are colored. The plot on the left is colored according to the ρ value, and the plot on the right is colored by the σ value. Notice that the correlated noise (ρ parameter) mainly affects the window-average fSNR and the single-bin SNR is mainly affected by the standard deviation parameter σ. With our focus on single-bin decoding, the simple Gaussian noise model was sufficient when generating synthetic datasets.

**Supplemental Table 1.** Two-sample T-tests for significance between tuning correlations. These p-values correspond to the bar plots in Figure 2c. Bolded entries indicate significance as any value below 0.01. BiH denotes H hand during the bimanual context, and UniH denotes H hand during the unimanual context.

| X-direction | UniL-BiL | UniR-UniL | BiR-BiL |
| --- | --- | --- | --- |
| UniR-BiR | **7.82 x 10<sup>-10</sup>** | **4.89 x 10<sup>-40</sup>** | **2.39 x 10<sup>-11</sup>** |
| UniL-BiL |  | **3.4 x 10<sup>-21</sup>** | 0.05 |
| UniR-UniL |  |  | **1.04 x 10<sup>-12</sup>** |

**Supplemental Table 1 | Two-sample T-tests for significance between tuning correlations.**
| Y-direction | UniL-BiL | UniR-UniL | BiR-BiL |
| --- | --- | --- | --- |
| UniR-BiR | **3.3 x 10<sup>-9</sup>** | **9.32 x 10<sup>-27</sup>** | **4.0 x 10<sup>-16</sup>** |
| UniL-BiL |  | 0.76 | **3.28 x 10<sup>-6</sup>** |
| UniR-UniL |  |  | **8.51 x 10<sup>-13</sup>** |

**Supplemental Table 2.** List of data collection sessions with participant t5. The trial day refers to the post-implant day.

| Date | Session # | Trial day | Description | Figure / Movie |
| --- | --- | --- | --- | --- |
| 06.02.2021 | 320 | 1750 | Cued unimanual and bimanual hand movement | Fig 1b,c<br>SFig 1b,c,e |
| 06.04.2021 | 321 | 1752 | Cued unimanual and bimanual hand movement (pilot data used to initially calibrate RNNs for closed-loop) | Fig 4a-c |
| 06.23.2021 | 324 | 1771 | Cued unimanual and bimanual hand movement (pilot data used to initially calibrate RNNs for closed-loop) | Fig 4a-c |
| 06.28.2021 | 325 | 1776 | Cued unimanual and bimanual hand movement, closed-loop two-cursor control | Fig 2b-d<br>Fig 4a<br>SFig 1d |
| 06.30.2021 | 326 | 1778 | Cued unimanual and bimanual hand movement, closed-loop two-cursor control | Fig 2b-d<br>Fig 4a<br>SMovie 1 |
| 07.12.2021 | 329 | 1790 | Cued unimanual and bimanual hand movement, closed-loop two-cursor control | Fig 4a |
| 07.14.2021 | 330 | 1792 | Cued unimanual and bimanual hand movement, closed-loop two-cursor control | Fig 2b-d<br>Fig 4a |
| 09.13.2021 | 336 | 1853 | Cued unimanual and bimanual hand movement, closed-loop two-cursor control (RNN vs. linear regression) | Fig 4c |
| 09.15.2021 | 337 | 1855 | Cued unimanual and bimanual hand movement, closed-loop two-cursor control (RNN vs. linear regression), 'Unimanual task' variant | Fig 4c<br>SMovie 3 |
| 09.27.2021 | 340 | 1867 | Cued unimanual and bimanual hand movement, closed-loop two-cursor control (augmented vs. non-augmented training data) | Fig 3e<br>Fig 5a-c<br>SFig 3a-c |
| 09.29.2021 | 341 | 1869 | Cued unimanual and bimanual hand movement, closed-loop two-cursor control (augmented vs. non-augmented training data) | Fig 3d-e<br>SMovie 4 |
| 10.11.2021 | 344 | 1881 | Cued unimanual and bimanual hand movement, closed-loop two-cursor control (sequential unimanual vs. simultaneous bimanual strategy) | Fig 2b-d<br>Fig 4a-b |
| 10.13.2021 | 345 | 1883 | Cued unimanual and bimanual hand movement, closed-loop two-cursor control (sequential unimanual vs. simultaneous bimanual strategy) | Fig 2b-d<br>Fig 4a-b<br>Fig 5a-c<br>SMovie 2 |

